# Scalable Extraction of Airway Mucins from Porcine Trachea

**DOI:** 10.1101/2024.10.08.617223

**Authors:** Sydney Yang, Elizabeth M. Engle, Allison Boboltz, Sahana Kumar, Alexa Stern, Gregg A. Duncan

## Abstract

Mucins are a major component of the innate defense system in the airways and their biological functions are important to consider in pulmonary disease research. However, the available mucus models for basic research relevant to the lung can be difficult to acquire in sufficient quantity to conduct such studies. Here, we present a new strategy to isolate airway mucins from pig trachea at the milligram to gram scale for use in pulmonary disease research. Using this protocol, we were able to isolate mucins with minimal DNA contamination consisting of ∼70% by weight protein. Compared to porcine gastric mucins extracted with the same procedure, the porcine tracheal extract possessed significantly greater O-linked glycoprotein (mucin) content. Particle tracking microrheology was used to evaluate the biophysical properties of porcine trachea mucins. We found porcine tracheal mucins formed a much tighter mesh network and possessed a significantly greater microviscosity compared to lab extracted porcine gastric mucins. In comparison to mucus harvested from human airway tissue cultures, we found porcine tracheal mucins also possessed a greater microviscosity suggesting these mucins can form into a gel-like material at physiological total solids concentrations. These studies establish an accessible means to isolate airway mucins from porcine trachea at large scale for use in pulmonary disease research.

## INTRODUCTION

Mucus lines the epithelium of mammalian organs including the stomach, eyes, respiratory tract, and reproductive tract.^1^ The mucus gel lining these tissues serves as a protective barrier against foreign particulates as well as providing lubrication and hydration.^2–4^ Together, mucins secreted at epithelial surfaces function to form the mucus gel network through electrostatic interactions, entanglement, and polymerization via disulfide bonding within cysteine rich domains of the mucin monomer.^2,5,6^ Mucus gels produced in distinct tissues possess unique biomolecular and biophysical properties depending on their precise functions. For example in the gut, a mucus gel layer exists with a loosely cross-linked, microbe-rich mucus layer overlying a more densely crosslinked, microbe-free mucus layer to physically separate the epithelium from the microbiome and other potentially pathogenic microbes.^7,8^ Prior studies have shown airway mucins can self-organize into strands and sheets to support mucociliary function and airway clearance.^9,10^ In addition, each mucin varies in its *O-*linked glycosylation pattern which contain different functional groups such as terminal sulfates, sialic acid, and fucose.^6,11^ Prior work has established these terminal functional groups directly impact the ability of airway mucins to neutralize viral pathogens and prevent infection.^12–14^ Thus, the structure and function of mucins is highly tailored to the tissue from which they arise.

Historically, mucins have been purified from animal tissues, such as bovine submaxillary mucins (BSM), porcine small intestinal mucins (PSIM), and porcine gastric mucins (PGM).^15^ These animal derived mucins are unique in their physicochemical properties, such as containing regionally specific mucins such as MUC2 and MUC5AC in the gastrointestinal tract.^7^ There are two commercially available mucins BSM and PGM, which will herein be referred to as BSM_C_ and PGM_C_ for clarification between lab-purified mucins and commercially purified mucins. While these commercial mucins are available in bulk making them convenient for use, a previous report has shown that these mucins contain significant amounts of DNA as well as other contaminants and may be partially degraded due to their processing.^16^ Mucins in the airway possess a unique composition, predominantly composed of MUC5B and MUC5AC,^2,5^ that is distinct from other mucosal tissues. As a result, previously reported lab extracted and commercial mucins do not provide suitable mucin/mucus models for airway disease research. Airway derived mucins have been primarily sourced from human patient samples (e.g. sputum^17^, mucus collected from endotracheal tubes^18,19^) or from human airway tissue cultures grown at air-liquid interface.^20–22^ This presents limitations in their broader use as patient-derived airway mucus is not widely available and tissue culture derived airway mucus is not easy to produce in large quantities. Motivated by this, we report a scalable extraction protocol, adapted from previous work,^23,24^ to isolate mucins from the porcine trachea. This establishes an approach to isolate airway mucins in an accessible and scalable manner which could be adopted broadly by researchers with interests in mucin-driven lung disease mechanisms.

## MATERIALS & METHODS

### PTM & PGM extraction from porcine tissue

Porcine trachea mucins (PTM) and porcine gastric mucins (PGM) were extracted based on a previously described protocol.^23^ Briefly, tissue samples were dissolved in 0.1 M NaOH overnigh. The porcine small intestine was filled with NaOH whereas the porcine trachea tissue was submerged in NaOH to allow for the solubilization of the mucus layer. Mucins were then removed from solution by lowering the pH to 4.0 with 1 M HCl (gel phase), then centrifuged at 3500 rpm for 20 minutes to allow for removal of the supernatant. The resulting pellet was resuspended in deionized water and the pH was adjusted to 8.0 to allow for resolubilization of the mucins (sol phase). The pH was altered to repeat the gel-sol cycle three times for further mucin extraction. Mucins were then purified by addition of DNase I (10 U/mL) at 21 C overnight. Following DNA removal, the gel-sol cycle was repeated 4 times through pH cycling and centrifugation for further mucin purification. Supernatant was then dialyzed (100 kDa) in deionized water for 72 hours, and then frozen at -80C overnight prior to lyophilization. The lyophilized extracted mucins were solubilized at 2% or 4% (w/v) in a physiological buffer containing 154 mM NaCl, 3 mM CaCl_2_, and 15 mM NaH_2_PO_4_, pH 7.4 prior to usage for further biochemical and biophysical analysis.

### BCi-NS1.1 cell culture & mucus collection

The immortalized BCi-NS1.1 human airway epithelial cell line was supplied by Ronald Crystal (Weill Cornell Medical College) and cultured as previously described.^25,26^ Briefly, BCi-NS1.1 cells were first expanded in a flask with Pneumacult-Ex Plus medium (no. 05040, StemCell Technologies) until confluent. Cells were then seeded (1 × 10^4^ cell/cm^2^) on rat tail collagen type 1-coated permeable Transwell membranes (12 mm; no. 38023, StemCell Technologies) until confluent. After confluence, only basal Pneumacult-ALI medium was provided in the basolateral compartment (no. 05001, StemCell Technologies) for 4 weeks to allow for polarization to occur at the air-liquid interface (ALI). Culturing at ALI allowed for the formation of an in vivo pseudostratified mucociliary epithelium. Mucus was collected through a 30 minute PBS wash, and concentrated using 100k MWCO amicon filters, and stored at -80C until usage.

### Mass DNA quantification assay

The mass DNA of extracted mucins was measured according to a previously established protocol.^24^ In brief, 30 μL of 20% (w/v) 3,5-diaminobenzoic acid was added to solubilized mucin samples, incubated at 60°C for 1 hour, and then the reaction stopped with addition of 1 mL of 1.76 M HCl. The reaction was stopped with the addition of 1 mL of 1.76 M HCl. The fluorescence intensity was measured at 390/530 nm (ex./emis.).

### Protein quantification assay

Protein concentration was quantified using a bicinchoninic acid (BCA) assay (no. 23225, ThermoFisher) as described by the manufacturer. In brief, a BCA working reagent was prepared, of which 200 μL was added to 15 μL of each sample, and then incubated for 30 minutes at room 37°C. Absorbance values of the samples were measured at 562 nm and compared to BCA standards.

### Relative O-linked glycoprotein content assay

O-linked glycoprotein (mucin) concentration was determined using a cyanoacetamide (CNA) reagent protocol as previously established.^24^ In brief, 200 µL of CNA was mixed with 1 mL of 0.15 M NaOH to create the CNA reagent. 60 µL of the CNA reagent were mixed with 50 µL of solubilized mucin samples (2% of 4% w/v) and incubated for 30 minutes at 100°C. The fluorescence intensity of the resulting samples was measured at 336/383 nm (ex./emis.) and compared to a serially diluted bovine submaxillary mucin (BSM; Sigma-Aldrich) standard curve.

### Sialic acid concentration assay

Sialic acid concentration was measured for extracted and commercial mucins by utilizing a modified Warren method centered on the thiobarbituric acid reaction (Sigma-Aldrich, MAK314) and following the manufacturer protocol. Briefly, bound sialic acid was hydrolyzed, then oxidized to form formylpyruvic acid which formed a measurable colored solution with the addition of thiobarbituric acid. The fluorescence of the colored solution was measured at 555/585 nm (ex./emis.) and compared to the sialic acid standard curve.

### Disulfide bond concentration assay

For extracted mucins the disulfide bond concentration was determined using a previously described protocol.^17^ In brief, 8 M Guanidine hydrochloric acid was added to 50 - 70 µL of sample for a final volume of 500 µL. 10% (v/v) of 500 mM iodoacetamide was added and samples sat at room temperature for 1 hour. 10% (v/v) of 1 M DTT was added, and samples incubated at 37°C for 2 hours. Samples were then filtered and buffer exchanged in a 7 KDa MWCO Zebra desalting column with 50 mM Tris-HCl (pH 8.0). As a standard, solutions of L-cysteine with concentration ranging from 0 µm – 5000 µM were prepared. Samples were diluted 1:1 with 2 mM monobromobimane in a flat black plate 96-well plate and incubated in the dark at room temperature for 15 minutes before reading the fluorescence at 395/490 nm (ex./emis.).

### Multiple particle tracking and microrheology analysis

Using a previously established protocol,^17^ nanoparticle (NP) probes were prepared for use in multiple particle tracking experiments by modifying 100 nm carboxylate-modified polystyrene NP (ThermoFisher) with a polyethylene glycol (PEG) coating to render these particles non-adheseive to mucus. We then constructed custom microscopy chamber consisting of a vacuum grease coated O-ring that was then filled with 20 µL of the mucin / mucus sample of interest and 1 µL of muco-inert NPs (∼0.002% w/v) before being sealed with a coverslip. Slides were then incubated at room temperature for 30 minutes in the dark prior to fluorescence imaging (Zeiss Confocal LSM 800, 63x water-immersion objective) to allow for sample equilibration. NP diffusion was imaged for 10 seconds at 33.3 frames per second and then tracked and analyzed using a custom MATLAB code that calculated the mean squared displacement, ⟨*MSD*(*τ*)⟩=⟨(*x*^2^+*y*^2^)⟩, for each particle. The MSD values were used to estimate the microrheological properties of the gel through the Stokes-Einstein relation, *G*(*s*) = 2*k*_*B*_*T*/(*πas*⟨Δ*r*^2^(*s*)⟩), where *k*_*B*_*T* is the thermal energy, *a* is the radius, and *s* is the complex Laplace frequency. The frequency-dependent complex modulus (G*) was calculated as, *G*^*^(*ω*)=*G*^*’*^(*ω*)+*G*^*”*^(*iω*) where *iw* is substituted for *s, i* is the complex number, and *w* is the frequency. The pore size (*ξ*) was estimated from the G’ using the following expression: *ξ*=(*k*_*B*_*T*/*G′*)^1/3^.

## RESULTS

### Biochemical properties of mucin extracted from porcine trachea

To better understand the composition of the resulting extracted mucin from porcine trachea (PTM_EX_), a series of biochemical assays were performed to assess DNA, total protein, *O-*linked glycoprotein (mucin), sialic acid, and disulfide bond content **(Fig. 1)**. For comparison, we conducted these analyses on commercial mucins (BSM_C_ and PGM_C_) as well as a lab extracted PGM harvested using the same protocol (PGM_EX_). The resulting measurements revealed that both extracted mucins, PTM_EX_ and PGM_EX_, contained minimal DNA, comparable to BSM_C_, and significantly lower DNA than PGM_C_ **(Fig. 1A)**. Similarly, both extracted mucins contained a mass protein of ∼70%, which was significantly higher than both BSM_C_ and PGM_C_ (**Fig. 1B)**. Measured *O-*linked glycoprotein (mucin) content normalized to that measured in BSM_C_, PTM_EX_ yielded a higher O-linked glycoprotein content compared to both PGM_C_ and PGM_EX_ **(Fig. 1C)**. Sialic acid concentration was similar in PGM_EX_ and PTM_EX_ **(Fig. 1D)**. In comparison to PGM_C_, the extracted mucins (PGM_EX_ and PTM_EX_) contained less sialic acid on average but these differences were not significantly different. The modified Warren assay which our sialic acid protocol employs has been reported to have difficulty hydrolysis of BSM_C_ mucins and as such, sialic acid content for BSM_C_ was not included in our results.^27^ Disulfide bond content was comparable between all mucin types **(Fig. 1E)**.

**Figure 1.**
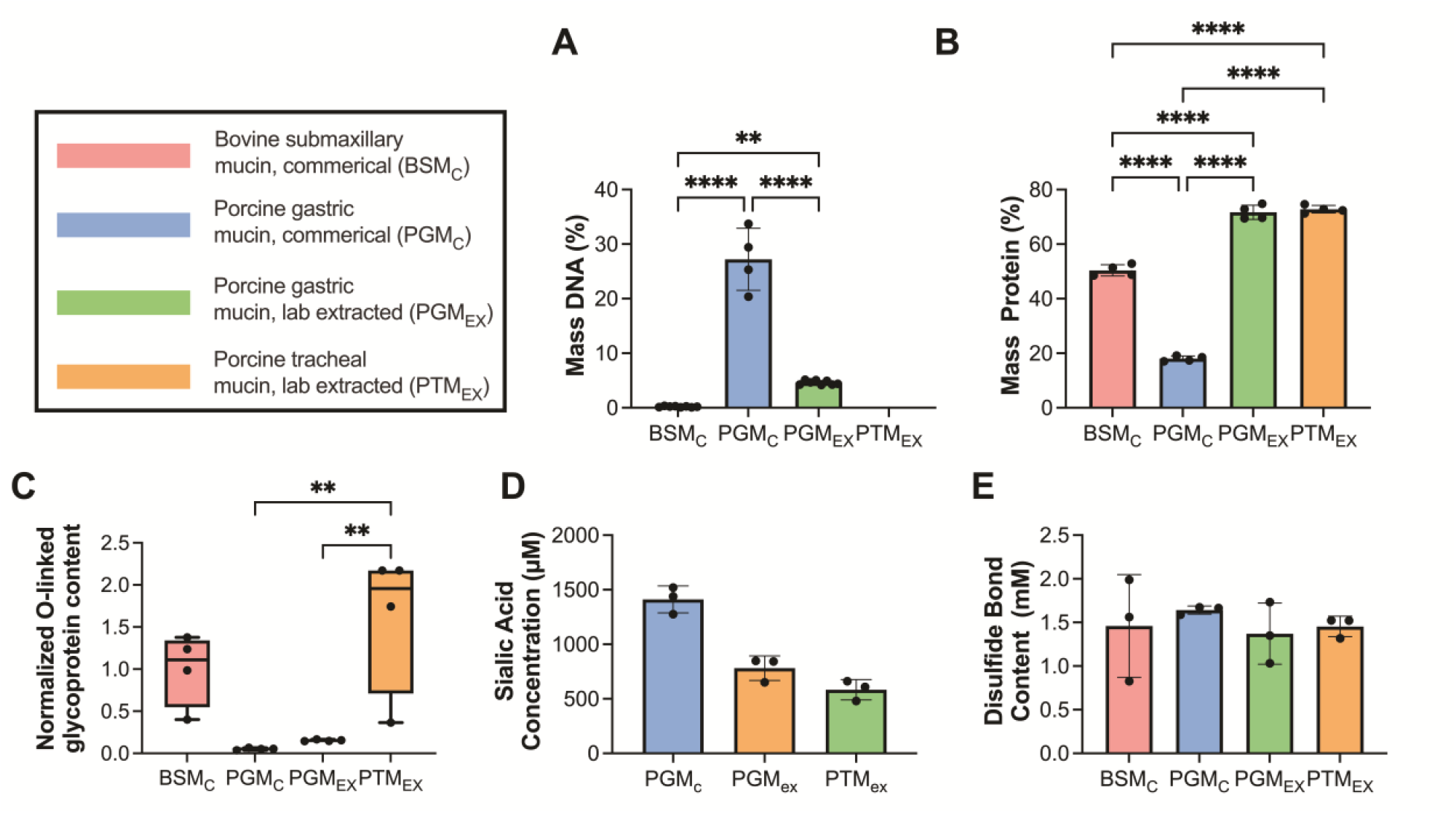
Biochemical characterization of mucins extracted from porcine trachea. (**A-E**) Commercially available BSM (BSM_C_), commercially available PGM type III (PGM_C_), laboratory extracted PGM (PGM_EX_), and laboratory extracted PGM (PGM_EX_) were characterized to determine (**A**) % mass DNA content, (**B**) % mass protein, (**C**) *O*-linked glycoprotein content normalized to BSM_C_, (**D**) sialic acid content, and (**E**) disulfide bond content. Bars indicate mean with error bars for standard deviations and individual measurements are shown as data points. Data set analyzes for statistical significance with Kruski- Wallis statistical test: *p<0.05, **p<0.01, and ****p<0.0001. Comparisons are not significant (p > 0.05) unless noted otherwise.

### Microrheological properties of mucin extracted from porcine trachea

We next used particle tracking microrheology to evaluate the viscoelastic properties of PGM_EX_ and PTM_EX_ prepared at physiological solids concentration of 2% and 4% w/v. Based on representative trajectories and measured mean squared displacement at a time scale of 1 s (MSD_1s_), NP were highly mobile within PGM_EX_ whereas NP diffusion appeared significantly constrained within PTM_EX_ **(Fig. 2A,B)**. The mean pore sizes for PGM_EX_ were ∼1.5–2.5 fold larger than PTM_EX_ where average pore sizes were ∼1 µm in PGM_EX_ and ∼500–750 nm in PTM_EX_ **(Fig. 2C)**. The mean microviscosity for PGM_EX_ was similar at both 2% and 4% w/v concentration. In comparison to PGM_EX_, PTM_EX_ possessed a 1.5 greater microviscosity at 2% w/v and a ∼13-fold greater microviscosity at 4% w/v PTM_EX_ **(Fig. 2D)**. Additional studies were conducted to compare the microrheological properties of PGM_C_ and PGM_EX_ where we found PGM_C_ formed a tighter network overall presumably due to the high DNA content in this mucin source **(Fig. S1)**. Similar studies were conducted comparing PTM_EX_ and BSM_C_ which showed PTM_EX_ possessed smaller pore sizes and greater microviscosity **(Fig. S2)**.

**Figure 2.**
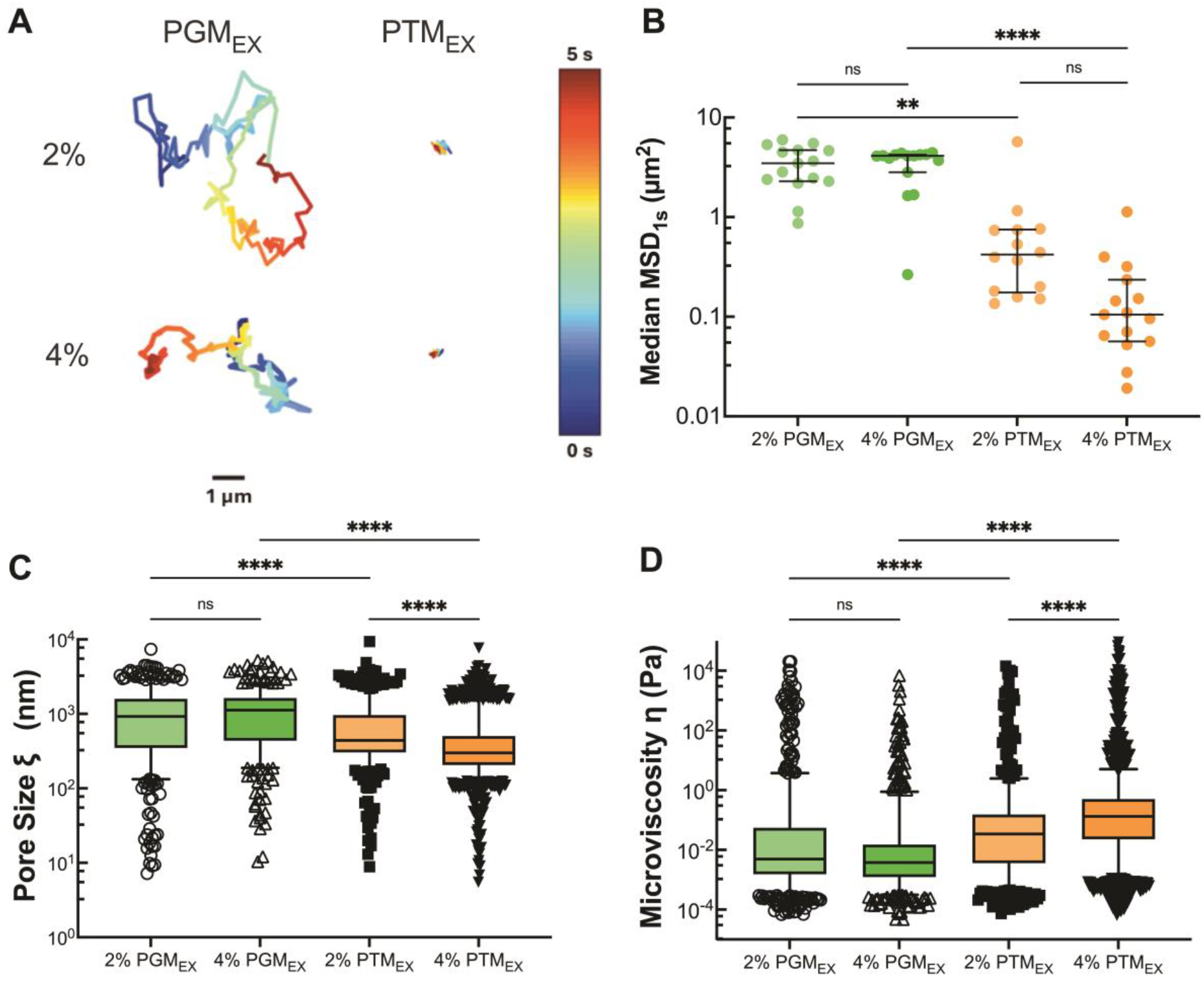
Microrheology of porcine gastric and tracheal mucins. (**A**) Representative trajectories of NP diffusion of 100 nm NP in 2% and 4% w/v PTM_EX_ and PGM_EX_. Trajectory colors change as a function of time with 0 s indicated by dark blue and 5 s indicated by dark red. Scale bar = 1 μm. (**B**) Calculated median MSD at a time scale of 1 second (MSD_1s_) for solubilized mucins. Each data point represents the median calculated MSD_1s_ in each video with at least 5 videos from 3 technical replicates. Black lines indicate interquartile range. (**C**) Estimated pore size (ξ) from NP diffusion. (**D**) Estimated microviscosity (η) from NP diffusion. Datasets in (**B**,**C**,**D**) analyzed with Kruski-Wallis test with Dunn’s test for multiple comparison: ns = not significant, **** p < 0.0001, ** p < 0.01.

To determine how the microrheological properties of PTM_EX_ compared to a traditionally used airway mucus model, we isolated airway mucus from differentiated human airway epithelial tissue cultures. Prior studies by our group have shown mucus collected from HAE cultures possess a total solids concentration of 2-4% w/v.^26^ Considering this, the microrheological properties of 2% and 4% w/v PTM_EX_ were compared to HAE mucus. NP trajectories in HAE mucus were observed to be comparable to 2% PTM_EX_ whereas diffusion was much more restricted in 4% PTM_EX_ **(Fig. 3A)**. Given the qualitative similarities, we quantitatively compared measured MSD_1s_, pore size, and microviscosity for 2% PTM_EX_ and HAE mucus. While similar in magnitude, we found a significant decrease in MSD_1s_ and pore size, indicative of tighter mesh network in 2% PTM_EX_ (**Fig. 3B,C**). PTM_EX_ also possessed a greater microviscosity compared to HAE mucus **(Fig. 3D)**.

**Figure 3.**
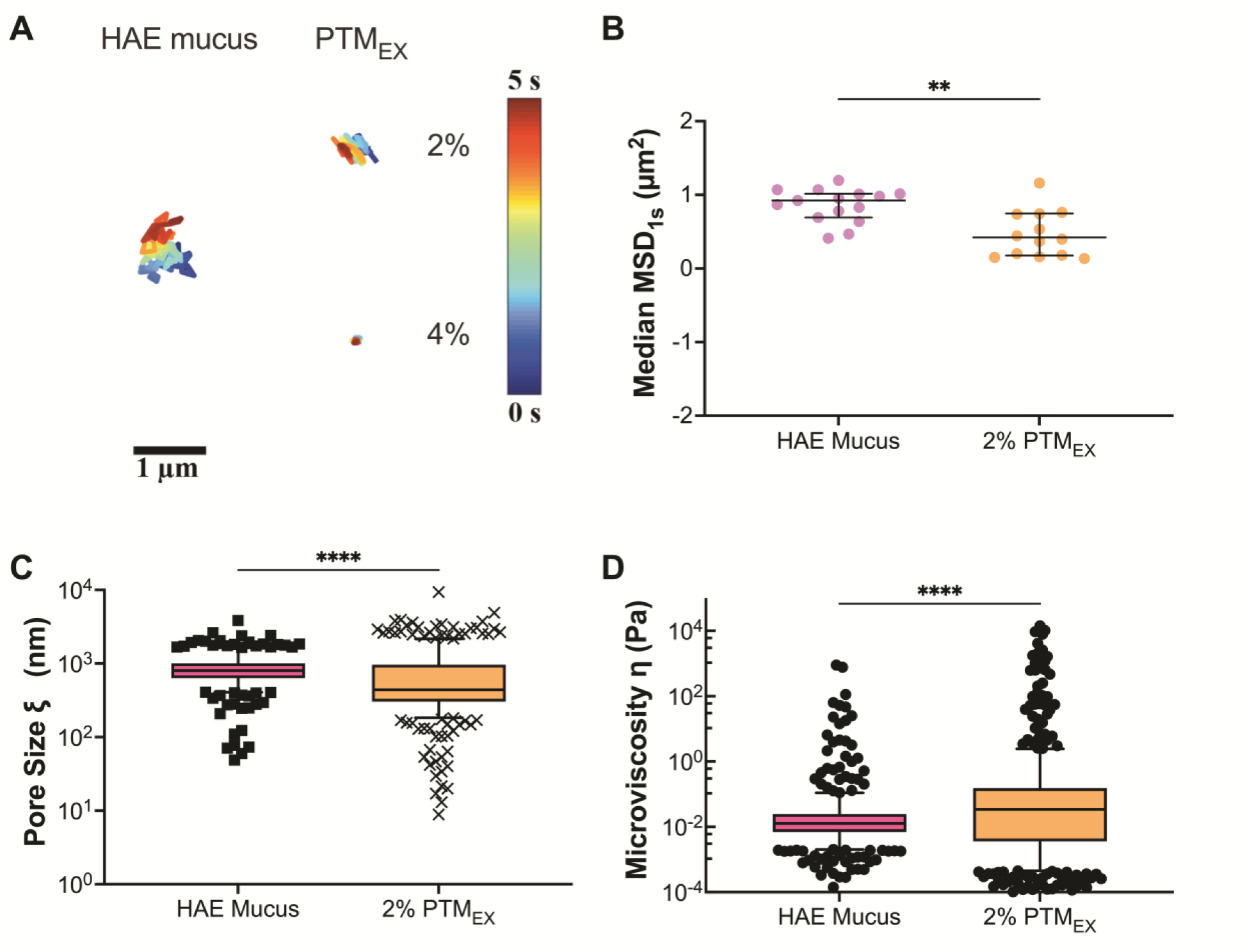
Microrheology of porcine tracheal mucins and mucus collected from human airway tissue cultures. measurements resulting from 100 nm nanoparticles in BSM_C_, and PTM_EX_. A: Representative trajectories for diffusion of 100 nm PS-NP in 2% solubilized mucins. Trajectory colors change as a function of time with 0 s indicated by dark blue and 5 s indicated by dark red. Scale bar = 1 μm. B: Calculated median MSD at a time scale of 1 second (MSD_1s_) for solubilized mucins. Each data point represents the median calculated MSD_1s_ in each video with at least 5 videos from 3 technical replicates. Black lines indicate interquartile range. C: Estimated pore size (ξ) from NP diffusion. D: Estimated microviscosity (η) from NP diffusion. Datasets in (B,C,D) analyzed with Mann-Whitney test: **** p < 0.0001, ** p < 0.01.

## DISCUSSION

We have developed a method for the extraction of porcine trachea mucins based on a previously established protocol by Sharma et al. Using a pH cycling method, airway mucins can be extracted at large scale (up to g) without the need for harsh chemical or enzymatic treatment. Similar to commercially available BSM_C_, the extracted mucins, PTM_EX_ and PGM_EX_, contained minimal DNA contaminants. PTM_EX_ contained the highest degree of extracted *O*-linked glycoproteins (mucin) compared to BSM_C_, PGM_C_, and PGM_EX_. We also quantified the content of terminal sialic acid glycans as these are relevant to many homeostatic and disease-associated processes in the airway. PTM_EX_ and PGM_EX_ both possessed lower sialic acid content compared to PGM_C_, although this is not necessarily indicative of issues with mucin glycoprotein integrity. Given PGM_EX_ possessed less sialic acid compared to its commercial counterpart, it is possible that our extraction method reduces free and/or mucin-associated sialic acid content.

Using particle tracking microrheology, we found PTM_EX_ forms a tighter network structure than PGM_EX_ with decreased diffusivity, reduced pore size, and increased microviscosity. This is most likely due to the larger fraction of O-linked glycoproteins (mucins) in PTM_EX_ which are the primary structural units of mucus gels. It should be noted mucus gels composed of PGM_EX_ are typically prepared at acidic pH.^28^ A comparison of pH-dependent behavior of PTM_EX_ and PGM_ex_ gel formation would be interesting to consider in future work. We also found PTM_EX_ and HAE culture derived mucus possessed similar network structure with a decreased pore size and increased overall microviscosity observed for PTM_EX_. These studies suggest that PTM_EX_ can form into a gel with similar physical structure to tissue culture-derived airway mucus and may be suitable as a model to study airway mucin function. Overall, this study establishes a new method for extraction of airway mucins with physiologically relevant biochemical and biophysical properties. This extraction method fills an important gap in the pulmonary research field to provide a more broadly accessible mucin source that is representative of airway mucus.

## LIMITATIONS OF THIS STUDY

We did not perform bulk rheological analyses to confirm formation of a gel using PTM_EX_ and we plan to perform these measurements in future work. It should be noted we did not measure the total solids content or other biochemical properties of the HAE mucus used in this study. A study comparing PTM_EX_ and HAE mucus with matched total solids concentration would be helpful in comparing these mucus sources in future work.

## ACKNOWLEDGEMENTS

This study was funded by the NIH (EB030834, HL160540, and HL160540-S1 awarded to S.Y.) and NSF GRFP (GRFP DGE1840340 awarded to S.Y.).

## AUTHOR CONTRIBUTIONS

S.Y. and A.B extracted and purified mucins. S.Y, E.M.E, A.B, and S.K performed all biochemical characterizations in Figure 1. E.M.E, S.K and A.S performed all microrheological tracking experiments and data analysis in Figure 2 and Figure 3. G.A.D. and S.Y. conceived and designed experiments. E.M.E and G.A.D. wrote the article. All authors reviewed and edited the article.

