## Supplemental Information for "Scalable Extraction of Airway Mucins from Porcine Trachea"

Sydney Yang[1]‡, Elizabeth M. Engle[1,2]‡, Allison Boboltz[1], Sahana Kumar[1,2], Alexa Stern[1], Gregg A. Duncan[1,2]
[1]Fischell Department of Bioengineering, [2]Molecular and Cellular Biology Program, University of Maryland, College Park, Maryland, United States of America, 20742

‡These authors contributed equally to this work.

### SUPPLEMENTAL INFORMATION

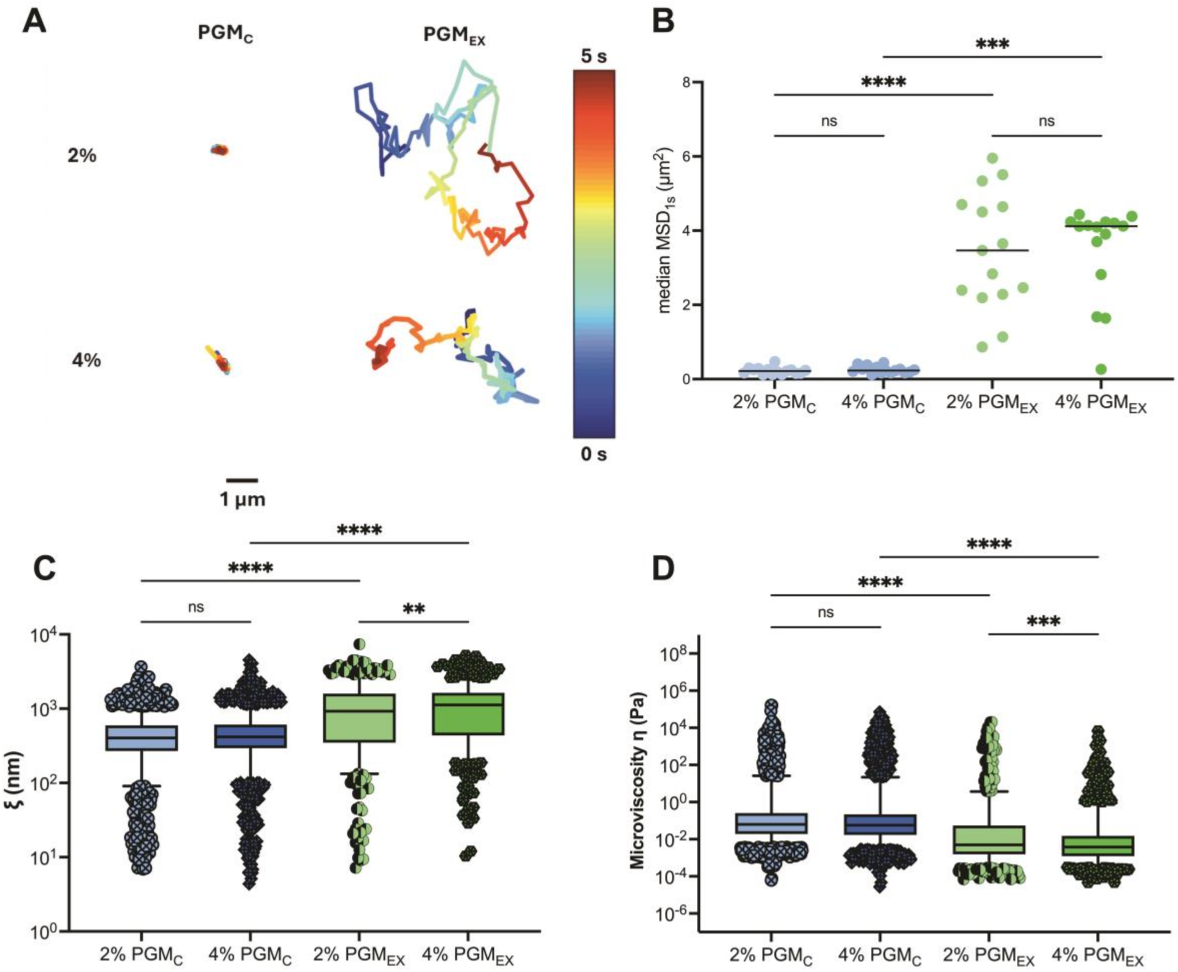


**Figure S1. Microrheology of commercial (PGM$_C$) and lab extracted (PGM$_{EX}$) porcine gastric mucins.** (**A**) Representative trajectories for diffusion of 100 nm NP in 2% and 4% solubilized mucins. Trajectory colors change as a function of time with 0 s indicated by dark blue and 5 s indicated by dark red. Scale bar = 1 μm. (**B**) Calculated median MSD at a time scale of 1 second (MSD$_{1s}$) for solubilized mucins. Each data point represents the median calculated MSD$_{1s}$ in each video with at least 5 videos from 3 technical replicates. Black lines indicate interquartile range. (**C**) Estimated pore size (ξ) from NP diffusion. (**D**) Estimated microviscosity (η) from NP diffusion. Datasets in (B,C,D) analyzed with Kruski-Wallis test with Dunn's test for multiple comparison: ns = not significant, **** $p < 0.0001$, ** $p < 0.01$.

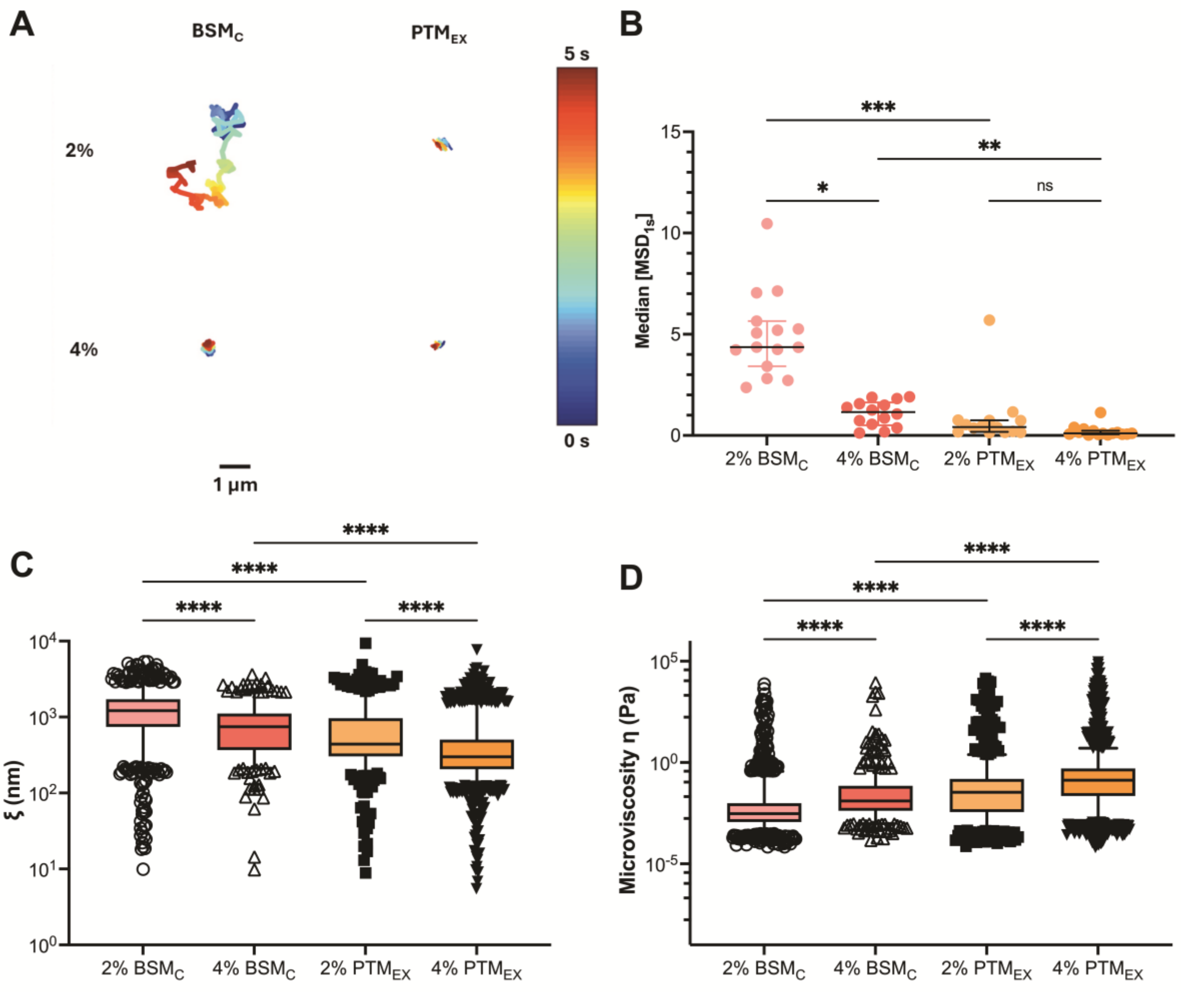


**Figure S2. Microrheology of bovine submaxillary ($BSM_C$) and porcine tracheal ($PTM_{EX}$) mucin. (A)** Representative trajectories for diffusion of 100 nm NP in 2% and 4% solubilized mucins. Trajectory colors change as a function of time with 0 s indicated by dark blue and 5 s indicated by dark red. Scale bar = 1 µm. (**B**) Calculated median MSD at a time scale of 1 second ($MSD_{1s}$) for solubilized mucins. Each data point represents the median calculated $MSD_{1s}$ in each video with at least 5 videos from 3 technical replicates. Black lines indicate interquartile range. (**C**) Estimated pore size (ξ) from NP diffusion. (**D**) Estimated microviscosity (η) from NP diffusion. Datasets in (B,C,D) analyzed with Kruski-Wallis test with Dunn's test for multiple comparison: ns = not significant, **** $p < 0.0001$, ** $p < 0.01$.
